# PHENOMS: Open-Source Software for Rust-Accelerated Hydrogen-Bond Analysis in Molecular Dynamics Simulations

**DOI:** 10.64898/2026.10.26.761768

**Authors:** Brandon Novy, Shu-Hang Lin, Rishabh Dey, Brenda Temple, Nicole Hajicek, Evgeniy V. Petrotchenko, Christoph H. Borchers, John Sondek, Konstantin I. Popov

**Affiliations:** UNC Eshelman School of Pharmacy, Center for Integrative Chemical Biology and Drug Discovery, Division of Chemical Biology and Medicinal Chemistry, University of North Carolina at Chapel Hill, Chapel Hill, NC 27599, USA; Segal Cancer Proteomics Centre, Lady Davis Institute, Jewish General Hospital, McGill University, Montreal, Quebec H3T 1E2, Canada; Department of Biochemistry and Biophysics, The University of North Carolina at Chapel Hill, Chapel Hill, NC 27599, USA; Department of Pharmacology, The University of North Carolina at Chapel Hill, Chapel Hill, NC 27599, USA; UNC Lineberger Comprehensive Cancer Center, The University of North Carolina at Chapel Hill, Chapel Hill, NC 27599, USA

## Abstract

Hydrogen-deuterium exchange mass spectrometry (HDX-MS) provides powerful insight into protein structure and dynamics, but interpretation of protection patterns at atomic resolution often requires complementary computational modeling. We present PHENOMS, an open-source Python package that integrates molecular dynamics trajectories with HDX-MS data to analyze backbone hydrogen-bond networks by tracking how individual hydrogen bonds and residue-residue contacts form, break, and reorganize throughout the simulations. This residue-level analysis can reveal local structural changes that may be missed by conventional global trajectory metrics. PHENOMS employs a multithreaded, Rust-accelerated computational engine that achieved an average 16-fold speedup (up to 100-fold) over widely used Python-based MD analysis packages in our benchmarks, while retaining individual hydrogen-bond resolution in every trajectory frame. We demonstrate PHENOMS through multiple retrospective case studies using published and in-house datasets, showing that it can recapitulate HDX-MS protection patterns, identify key residues that drive coordinated structural changes, and benchmark the computational efficiency of the approach. With automated quality-control workflows, PHENOMS provides a framework for integrating hydrogen-bond analysis with HDX-MS data or serves as a standalone, high-throughput tool for analyzing MD trajectories for broader applications in protein engineering, drug discovery, and structural biology.

## 1 INTRODUCTION

Hydrogen-deuterium exchange mass spectrometry (HDX-MS) is a powerful technique in structural biology that provides information on protein secondary structure.^1,2^ Differential HDX-MS can experimentally compare the hydrogen-bonding status of backbone amides across multiple protein states, providing information on conformational transitions and stable secondary structure elements.^3,4^ Combining differential HDX-MS with molecular dynamics (MD) simulations can extend the interpretation of peptide-level HDX-MS data to the level of individual hydrogen bonds,^5–10^ thereby enabling an atomistic description of protein conformational changes, including those induced by ligand binding.^7,11,12^ Specifically, we have shown that changes in hydrogen-bond occupancies and lifetimes in MD simulation trajectories correlate well with experimentally measured changes in HDX protections.^11,12^

Although modern AI-based structure prediction and co-folding methods have advanced the field, they may not fully capture subtle structural perturbations induced by ligand binding or single-point mutations, and their predictions remain less reliable for intrinsically disordered or highly dynamic protein regions.^13–15^ Integrating differential HDX-MS with physics-based MD simulations therefore provides a powerful complementary approach for resolving these dynamic and mechanistic changes at near-atomic resolution.

Here, we present PHENOMS (**P**ython-based **H**ydrogen-bond **E**valuation, **N**etwork analysis, and **O**ccupancy mapping from **M**olecular **S**imulations.), an open-source software package that unifies established and newly developed methods^11,12,16^ for automated analysis of backbone hydrogen-bond networks from MD trajectories, implemented and optimized for robust and efficient execution. Beyond its application to interpreting HDX-MS data, PHENOMS provides a general mode for analyzing all hydrogen bonds and their networks in MD simulations. Its memory-efficient data processing enables the analysis of large systems and long trajectories. The framework automatically standardizes residue numbering, performs automated quality assessment of trajectory replicates and parameter consistency to evaluate convergence across states, and generates graph-based visualizations of hydrogen-bond donor-acceptor network reorganization. In our multi-threaded benchmark, PHENOMS, powered by Rust-accelerated engine, identifies hydrogen bonds 7-to 22-fold faster than MDAnalysis^17^ and up to ∼115-fold faster than MDTraj^18^, two widely used packages for MD trajectory analysis. We demonstrate the application of PHENOMS workflows for analyzing protein mutations using the EGFR Δ747–749 mutant,^19,20^ a clinically significant exon 19 deletion and a primary therapeutic target in oncology.^21^ PHENOMS successfully identifies the EGFR L858R hotspot mutation site, one of the most abundant activating mutations in non-small cell lung cancer (NSCLC), and captures protein dynamic behavior that directly corroborates published HDX-MS observations.^20,22^ PHENOMS is available through PyPI and includes validated functions and a comprehensive unit test suite, supporting reproducible, high-throughput analysis for protein engineering and drug discovery applications.

## 2 DESIGN AND IMPLEMENTATION

PHENOMS is an open-source Python package developed around the hydrogen-bond analysis of molecular dynamics (MD) trajectories. Initially designed to facilitate atomic-level interpretation of hydrogen-deuterium exchange mass spectrometry (HDX-MS) data, PHENOMS provides a robust framework for quantifying local interaction networks that are often not resolved by standard global metrics, such as root-mean-square deviation (RMSD) and root-mean-square fluctuation (RMSF). The software implements an object-oriented architecture centered on the ‘SimulationSet’ and ‘ComparisonSet’ classes, enabling automated handling of simulation replicates, state-to-state comparisons (e.g., wild-type vs. mutant), and the generation of publication-quality visualizations.

### Trajectory preparation and standardization

For user convenience, PHENOMS uses the widely adopted MDTraj library for initial loading and preparation of trajectories and topologies. To ensure consistency across diverse simulation engines, PHENOMS includes automated preprocessing utilities to standardize raw data, as well as scripts to handle periodic boundary condition removal, rotational and translational fitting, and an option to reduce the number of trajectory snapshots before analysis. To facilitate comparative studies of protein variants containing substitutions, insertions, or deletions, we implemented renumbering tools based on sequence alignment. These tools map residue indices to a common reference numbering scheme, enabling consistent comparison of differential protection metrics across variants.

### Accelerated Engine for the Hydrogen-Bond Detection

Hydrogen-bond identification is performed using Baker–Hubbard geometric criteria (hydrogen– acceptor distance < 2.5 Å and donor–hydrogen–acceptor angle > 120°), that are evaluated independently for every MD frame.^23^ For each identified hydrogen bond, PHENOMS computes a residue-pair occupancy (the fraction of simulation frames in which the bond is present) and a bond persistence score (the longest continuous run of frames over which the bond remains formed), expanding methodologies described in our previous publications.^11,12,24^ PHENOMS supports two primary analysis modes: a “Backbone Mode” focused on N—O bonding relevant to HDX-MS protection, and an “All-Bond Mode” for broader global interaction profiling. Computational efficiency is achieved through a multi-threaded Rust extension (‘phenoms_hbond_rs’), which performs kernel execution over flattened coordinate arrays. In our multi-threaded benchmarks, this implementation identified hydrogen bonds 7-to 22-fold faster than MDAnalysis and up to ∼115-fold faster than MDTraj, two established MD trajectory-analysis packages (Table 1). The large tables of per-frame bonds generated during analysis are organized and combined using the Polars data-processing library, allowing PHENOMS to efficiently analyze long simulation trajectories while minimizing manual data preparation.

**Table 1.** Docker-Based Multi-Threaded Benchmark of All H-Bond Identification for PLCγ2 MD Simulation conducted using the Amber MD Engine (19,174 atoms total). All benchmarks were executed within a Docker container using the phenoms-bench:4cpu image (available on GitHub; ∼5.2 GB; it can also be rebuilt from the Dockerfile provided in the PHENOMS repository), with a CPU quota limited to 4 vCPUs. Threading environment variables were fixed at 4, and each method was run with four worker threads in accordance with the respective software documentation. MDTraj and the Rust-based implementation both employ the native Baker–Hubbard algorithm for hydrogen-bond detection (hydrogen–acceptor distance < 2.5 Å donor–hydrogen–acceptor angle > 120°). MDAnalysis HydrogenBondAnalysis was run with its default geometric criteria, which differ from Baker–Hubbard, using the following selections and cutoffs: donors (protein and (element N or element O or element S)), hydrogens (protein and element H), and acceptors (protein and (element N or element O or element S)); a donor–hydrogen cutoff of 1.2 Å (used only to assign covalent donor–hydrogen pairs); a donor–acceptor distance cutoff of 3.0 Å; a donor–hydrogen–acceptor angle cutoff of 150°; and update_selections=False. Total columns report the hydrogen bonds returned by each package: per-frame instances for PHENOMS and MDAnalysis, and unique bonds passing MDTraj’s default 10% frequency filter for MDTraj.

| Window | Residues | Frames | Rust (s) | MDTraj (s) | MDAnalysis (s) | MDTraj/Rust | MDA/Rust | Rust_Total | MDTraj_Total | MDAnalysis_Total |
| --- | --- | --- | --- | --- | --- | --- | --- | --- | --- | --- |
| Full | 1177 | 100 | 0.343 | 39.533 | 3.742 | 115.275 | 10.912 | 115958 | 1911 | 74249 |
| Trunc1_1_1077 | 1077 | 100 | 0.370 | 11.799 | 4.103 | 31.863 | 11.080 | 103283 | 1701 | 65831 |
| Trunc2_1_977 | 977 | 100 | 0.271 | 6.718 | 1.912 | 24.763 | 7.047 | 93915 | 1556 | 59981 |
| Trunc3_1_877 | 877 | 100 | 0.191 | 5.814 | 1.840 | 30.439 | 9.634 | 81749 | 1340 | 52198 |
| Trunc4_1_777 | 777 | 100 | 0.153 | 2.218 | 1.390 | 14.503 | 9.085 | 70947 | 1173 | 44960 |
| Trunc5_1_677 | 677 | 100 | 0.133 | 2.112 | 1.240 | 15.857 | 9.309 | 63042 | 1036 | 39962 |
| Trunc6_1_577 | 577 | 100 | 0.091 | 1.092 | 1.328 | 12.057 | 14.660 | 53625 | 891 | 33780 |
| Trunc7_1_477 | 477 | 100 | 0.082 | 0.765 | 1.002 | 9.306 | 12.196 | 44744 | 746 | 28139 |
| Trunc8_1_377 | 377 | 100 | 0.040 | 0.508 | 0.785 | 12.574 | 19.441 | 36413 | 601 | 22757 |
| Trunc9_1_277 | 277 | 100 | 0.021 | 0.311 | 0.475 | 14.565 | 22.233 | 26871 | 445 | 17046 |

### Replicate Aggregation and Quality Control

The ‘SimulationSet’ model performs rigorous statistical analysis by aligning hydrogen-bond labels across replicate runs and generating pivoted occupancy matrices. These matrices organize the data for direct comparison, with each row representing an individual hydrogen bond and each column corresponding to a simulation frame or replicate. This format allows users to compare how often and how consistently each bond forms across runs. To ensure the reliability of downstream comparative analysis, PHENOMS incorporates an integrated quality control module. To identify trajectories that may not have reached equilibrium, this module performs an initial quality control check based on RMSD stability. The module takes the final 20% of the trajectory and divides it into two equal, consecutive blocks of frames. It calculates the average RMSD within each block and flags a potential shift if the two averages differ by more than 5%. Additionally, it fits a straight line to the RMSD values within the last block and flags trajectory drift if the absolute value of the slope exceeds a small threshold (1×10^−3^ Å per frame), indicating a persistent upward or downward trend in the RMSD. Together, these checks flag simulations that may not have reached equilibrium or may still exhibit structural drift, allowing users to consider these potential limitations when interpreting the analysis results. It can also verify that key MD simulation parameters (e.g. content of gromacs^25^ .mdp files), such as temperature and force-field settings, are consistent across input sets. These checks prevent the silent propagation of errors derived from unconverged or inconsistent simulation replicates.

### Differential Analysis and Network Topology

The ‘ComparisonSet’ class compares hydrogen-bond networks between two sets of simulations. This analysis can reveal differences between protein states, such as wild type and mutant proteins or apo and ligand-bound forms. The class generates a common list of all hydrogen bonds observed in either set, including those shared between conditions and those unique to one condition, and uses this list to quantify differences in hydrogen-bond behavior at the residue level. Beyond individual bond metrics, PHENOMS utilizes graph-based analysis to transform bond-level data to represent residue-level connectivity. This helps visualize how hydrogen-bond networks reorganize between simulations and identify local subnetworks of interacting residues that may stabilize distinct conformational states or participate in coordinated structural changes. Based on the hydrogen-bond network analysis, PHENOMS calculates residue-level scores that serve as proxies for changes in protection against hydrogen exchange. This scoring assesses changes in hydrogen bonds associated with individual residues to estimate whether their backbone amide hydrogens are more or less likely to be protected from exchange in one simulation condition relative to another, allowing comparison with experimental differential HDX-MS measurements. These scores can be exported directly into PDB B-factors for three-dimensional structural annotation and visualization in molecular graphics software such as PyMOL^26^ or Chimera^27^.

## 3 RESULTS

### Biological Benchmark

To demonstrate the utility of PHENOMS, we applied it to a well-studied and highly dynamic protein, the epidermal growth factor receptor (EGFR). Previous studies have used HDX-MS to characterize wild-type (WT) EGFR and several clinically relevant variants. Analysis of atomistic MD simulations of WT EGFR and the clinically relevant Δ747-749 variant using PHENOMS revealed global changes in protein dynamics and several localized regions with altered hydrogen-bond interactions (**Fig. 2**). Previous HDX-MS studies by van Alderwerelt van Rosenburgh et al.^20^ demonstrated that exon 19 mutations can alter hydrogen exchange and protein dynamics in EGFR. Consistent with these previous findings, our automated report from PHENOMS indicated the A-loop region is among the regions exhibiting the largest increases in conformational dynamics. This region contains the clinically important L858 mutation site.

**Figure 1.**
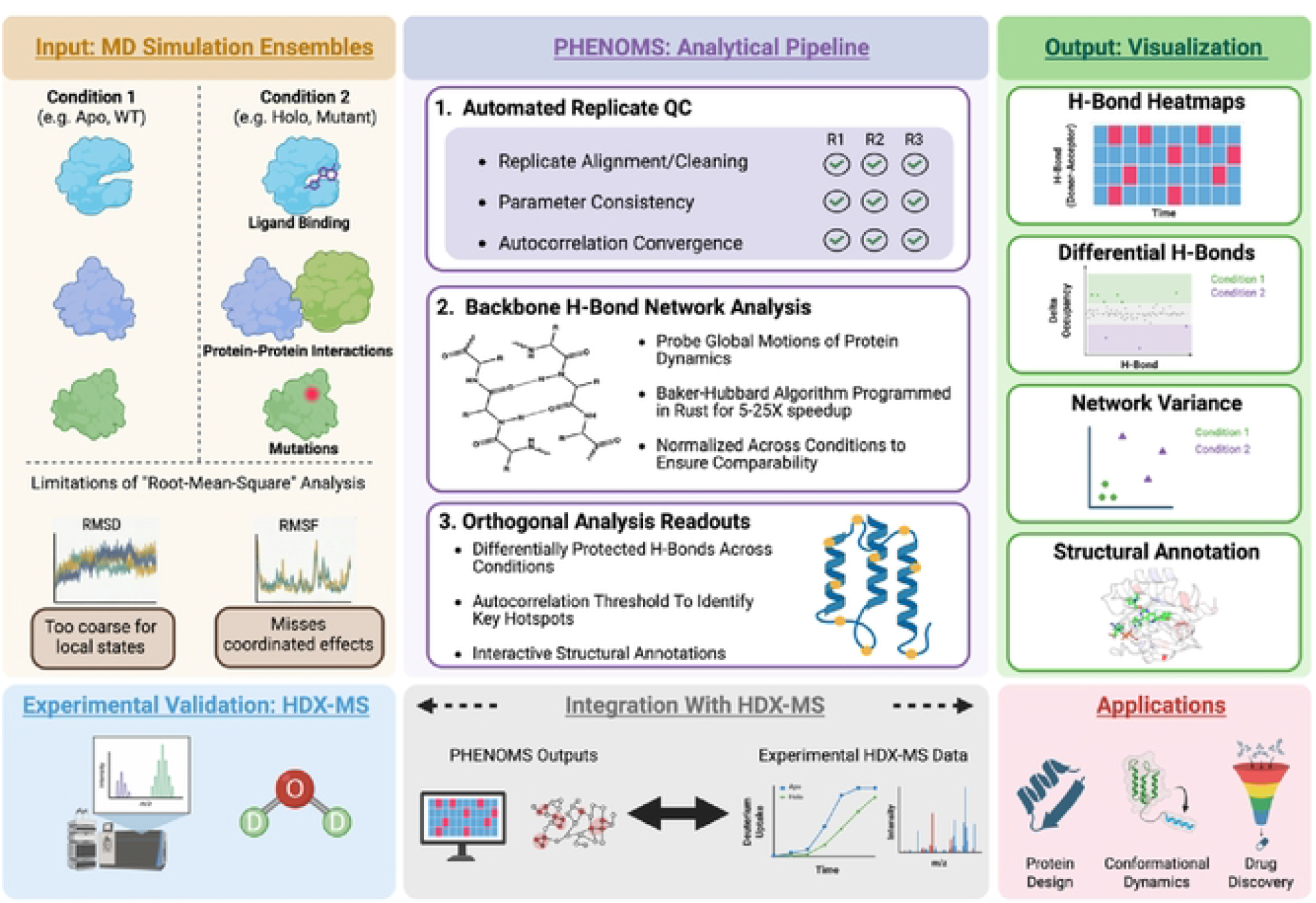
PHENOMS Python package suite overview. PHENOMS takes replicate MD trajectories from multiple protein states as input and generates multiple orthogonal, quantitative metrics describing protein structure and dynamics. These readouts can be integrated with HDX-MS data to experimentally validate and characterize structural and dynamic differences between the input states. The package implements an optimized workflow featuring: (1) automated replicate quality control to assess internal consistency across simulations: (2) backbone hydrogen-bond (H-bond) network analysis that extends beyond conventional global metrics such as RMSD and RMSF to resolve residue-level and secondary structure-specific dynamics: and (3) graph-based visualization and data export for multi-state comparisons, community analysis, and structural mapping through annotation of PDB B-factor fields. Optional integration with experimental HDX-MS measurements enables direct comparison between simulated and experimentally observed changes, providing a validated framework for investigating protein dynamics and supporting applications in protein engineering and drug discovery.

**Figure 2.**
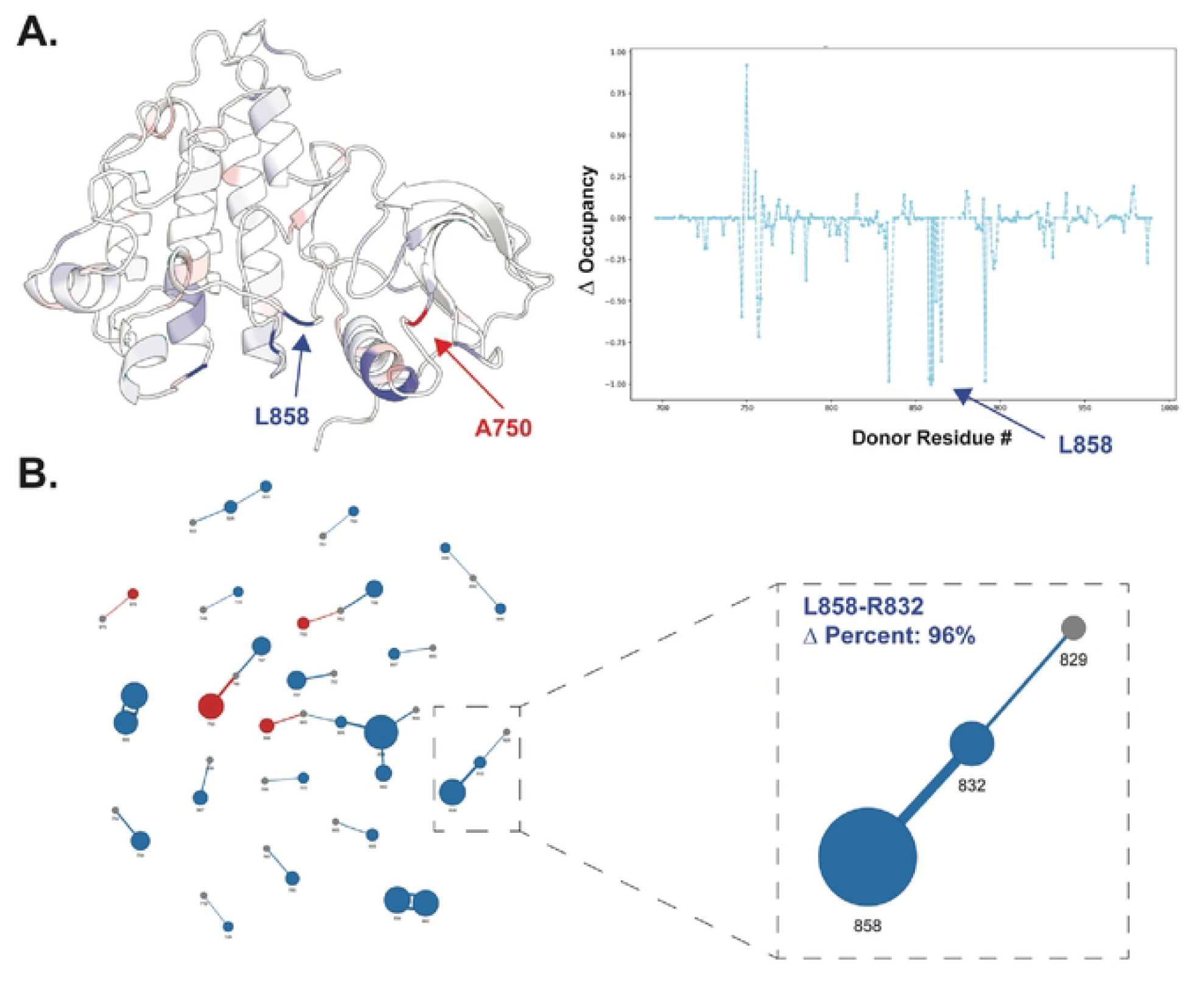
PHENOMS proof-of-concept validation using EGFR WT vs. Δ747-749. Analysis of all-atom MD simulations perfonned in triplicate, with nearly 1 µs of total sampling, revealed hydrogen-bond occupancy patterns consistent with previously reported HDX-MS protection data. (A) Structure of WT EGFR kinase domain (PDB: 1M17) annotated with average Changes in backbone hydrogen-bond occupancy between triplicate simulation conditions. The occupancy line plot highlights the strongest hydrogen-bond donor signals, with absent hydrogen bonds assigned zero occupancy. Analysis of temporal correlations in hydrogen-bond occupancy identifies L858 as a hotspot of altered hydrogen-bond dynamics. Notably, this residue is the site of the clinically important activating L858R mutation in NSCLC. (B) Analysis and visualization of hydrogen-bond networks as graphs, with residues represented by nodes and donor-acceptor interactions by edges. The highlighted region shows the interaction between R832 and L858, as displayed in the interactive HTML output of PHENOMS. Blue edges indicate hydrogen bonds stabilized in WT EGFR, whereas red edges indicate those stabilized in the Δ747-749 mutant (PDB: 7TVD).

### Algorithmic Efficiency Benchmark

To quantify the performance gains of the Rust-accelerated engine, we benchmarked the computationally demanding task of identifying hydrogen-bond networks in all-bond mode using phospholipase C gamma 2 (PLCγ2), a large protein comprising 19,174 atoms that we have previously studied.^28–30^ We compared PHENOMS against two widely used Python packages for MD trajectory analysis, MDTraj and MDAnalysis, using an identical four-thread configuration within a shared Docker environment (**Table 1**). For the full 1,177-residue PLCγ2 protein, the Rust engine completed the analysis of 100 trajectory frames in 0.34 s, compared with 3.74 s for MDAnalysis and 39.5 s for MDTraj, corresponding to approximately 11-fold and 115-fold speedups, respectively. This performance advantage held across progressively truncated PLCγ2 systems ranging from 1,177 to 277 residues, with PHENOMS achieving approximately 7-to 22-fold speedups over MDAnalysis in the 100-frame benchmarks (**Table 1**). For longer trajectories of 1,000 frames, PHENOMS maintained approximately 4-to 14-fold speedups over MDAnalysis, with the largest relative gains observed for smaller systems (**Table S1**). MDTraj was excluded from the 1,000-frame benchmarks due to its memory demanding requirements within the benchmarking virtual machine. PHENOMS and MDTraj use identical Baker-Hubbard geometric criteria for hydrogen-bond identification, allowing a more direct comparison of their computational performance. In contrast, MDAnalysis was evaluated using its default hydrogen-bond definition (donor–acceptor distance < 3.0 Å and donor–hydrogen–acceptor angle > 150°), representing typical out-of-the-box usage rather than a geometry-matched comparison. PHENOMS reported more hydrogen bonds than either package (**Table 1**). The higher number of hydrogen bonds reported by PHENOMS relative to MDTraj reflects the latter’s default filtering, which excludes bonds detected in fewer than 10% of trajectory frames. Relative to MDAnalysis, the difference largely reflects its stricter 150° angular cutoff. Retaining the complete record of hydrogen bonds in every trajectory frame is a deliberate feature of PHENOMS, as transient and low-occupancy interactions may contribute to changes in hydrogen-bond networks between protein states. These interactions are captured by the occupancy and persistence metrics calculated by PHENOMS, which maintains this level of detail while outperforming both packages in our benchmarks. The Docker environment used for benchmarking is distributed with PHENOMS, allowing users to reproduce or extend these comparisons on their own hardware.

## 4 CONCLUSIONS

PHENOMS provides a high-throughput, end-to-end framework for describing protein dynamics through the analysis of hydrogen-bond networks. By moving beyond conventional metrics like RMSD and RMSF, the package enables researchers to quantify the relational context of intramolecular interactions via residue-pair occupancy, bond persistence, and graph-based visualizations of donor-acceptor rewiring. The integration of a Rust-accelerated engine ensures that these complex network analyses remain computationally feasible for long-scale and large-size MD simulations, delivering the performance required for modern drug discovery and protein engineering workflows.

The development of PHENOMS is a direct response to the ongoing “reproducibility crisis” currently facing the life sciences. Recent research indicates that fewer than half of published Jupyter notebooks can be successfully executed, a mere 3% can fully reproduce original study results with functional dependencies,^**31**^ and only 51% of open-source packages can be installed with relative ease.^**32**^ To address these systemic issues, we have prioritized deployment stability and rigorous validation. PHENOMS is available via PyPI for effortless installation, supported by a comprehensive unit test suite and a Docker-based workflow to ensure consistent execution across different environments.

Ultimately, PHENOMS empowers the structural biology community to bridge the gap between atomistic simulations and experimental observations. By providing a mechanistically interpretable API for quantifying changes in protein dynamics, secondary-structure rearrangements, and associated hotspot residues, PHENOMS can help identify novel therapeutic targets and improve our understanding of protein function. Whether used as a companion tool for HDX-MS interpretation or as a standalone platform for MD analysis, PHENOMS provides the rigor and performance needed to translate raw data from MD trajectories into biological insight.

## 5 AVAILABILITY AND FUTURE DIRECTIONS

PHENOMS is implemented in Python and Rust and is openly developed on GitHub (https://github.com/Popov-Lab-UNC/PHENOMS), with tagged releases distributed via PyPI for installation with pip or uv (“pip install phenoms” or “uv add phenoms”). The package exposes both a command-line interface and a Python API, allowing it to be run as a standalone analysis tool or scripted directly into existing MD workflows. Comprehensive documentation, including installation instructions, tutorials, and a full API reference, is built with Sphinx and hosted online at https://popov-lab-unc.github.io/PHENOMS/. The package emphasizes reproducibility through a comprehensive suite of unit tests and a Docker-based benchmarking workflow that allows users to reproduce the performance results reported here with a single command. Typical PHENOMS applications include analyzing the effects of amino acid substitutions on secondary-structure stability, investigating conformational changes induced by ligand binding, and identifying networks of interacting residues associated with protein conformational stability. These capabilities support applications in protein engineering and drug discovery.

## DATA AVAILABILITY

The PHENOMS source code is freely available under the MIT license at https://github.com/Popov-Lab-UNC/PHENOMS and from PyPI (https://pypi.org/project/phenoms/). Documentation is available at https://popov-lab-unc.github.io/PHENOMS/. MD simulation data for EGFR WT and Δ747–749 and the PLCγ2 benchmark data are available on Zenodo (https://doi.org/10.5281/zenodo.22879722).

## Supplementary Data

All relevant data analyzed in this manuscript has been deposited on Zenodo at: https://zenodo.org/records/22879722 (10.5281/zenodo.22879722). Instructions for downloading and working with this data can be found in the documentation for PHENOMS at: https://popov-lab-unc.github.io/PHENOMS/data.html.

**Table S1.** Docker-based multi-threaded benchmark of all hydrogen-bond identification for the PLCγ2 MD simulation (19,174 atoms) over 1,000 frames, comparing PHENOMS with MDAnalysis. MDTraj was excluded owing to its memory requirements within the benchmarking VM.

## Conflict of Interest

The authors declare no competing financial interest.

## Funding

The authors gratefully acknowledge support from the NIH Biophysics Training Grant (T32GM148376-01A1). This research was partially funded by grants from the National Institutes of Health (R01AG098153 and R35GM149299).

## Notes

### Competing Interest Statement

The authors have declared no competing interest.

